# Distribution of midleg campaniform sensilla in Hemiptera and their association with vibrational communication

**DOI:** 10.64898/2026.10.11.758358

**Authors:** Brian Saltin, Leonidas-Romanos Davranoglou

## Abstract

Campaniform sensilla are cuticular strain receptors that contribute to locomotion, postural control and vibration detection in insects. Their distribution in Hemiptera remains incompletely documented, limiting comparative and experimental investigation of their possible role in vibrational communication. We used scanning electron microscopy to map externally recognisable campaniform sensilla on the midlegs of a phylogenetically broad sample of Hemiptera, with an emphasis on Heteroptera. We recorded sensilla number and location by leg segment, distinguished between confidently identified sensilla and candidate structures, and compared counts between taxa with documented vibrational communication and those without a published record. We found widespread high concentration of campaniform sensilla across Heteroptera, although the distribution of sensilla on other leg segments was more variable. The sampled Pentatomoidea Trichophora had particularly high counts, and the exploratory comparison indicated higher counts in taxa with documented vibroacoustic communication. This association requires cautious interpretation because communication is incompletely documented, sampling is limited, and sensilla number may also reflect shared ancestry, leg mechanics and other sensory functions. Our survey provides a comparative anatomical basis for testing how campaniform sensilla contribute to vibration perception in Hemiptera.

## Introduction

Arthropods possess a diverse array of mechanoreceptive organs, including external sensors like hair sensilla that respond to air movement, and internal ones such as scolopidial sensilla, which are especially sensitive to vibration (Keil, 1997). Among the most widespread external mechanoreceptors in arthropods are the dome-shaped campaniform sensilla (CS), which detect cuticular deformation and strain (Keil, 1997). These sensilla are typically located in mechanically active regions, such as leg joints (Mendes et al. 2014), wings (Ai et al., 2010; Dickerson et al., 2021), and mouthparts (Brożek and Chłond, 2010; Parveen et al., 2015; Taszakowski et al., 2019; Wang and Dai, 2017), and may occur individually or in arrays.

Campaniform sensilla are versatile general mechanosensors (Dickerson et al., 2021), informing a multitude of insect behaviours. In Diptera, they participate in haltere-mediated flight control (Dickinson, 1999; Pringle, 1948; Simmons, 2024; Smith, 1969; Yarger and Fox, 2018), and at the same time they have also been implicated directly or indirectly in vibration reception in other insects (Čokl and Amon, 1980; Kühne, 1982; Markl, 1970). Furthermore, their long-established role as load sensors is considered crucial to insect locomotion (Akay et al., 2001; Büschges et al., 2008; Harris et al., 2022; Zill et al., 2004, 2014, 2011, 2010, 2009, 1999; Zill and Moran, 1981).

The directional sensitivity and specific orientation of CS on appendages such as legs, wings, or halteres allow them to contribute to posture control, coordinated locomotion, and proprioceptive feedback loops (Dickerson et al., 2021; Lakes-Harlan and Strauß, 2014; Strauß, 2020).

The distribution of CS in Hemiptera is poorly documented, particularly on the legs and abdomen, which may participate in the reception of mechanical signals. Within Heteroptera, a suborder comprising approximately 40,000 species, a trochanteral field of CS is commonly termed as the trochanteral organ. This structure has been examined primarily for taxonomic purposes in the infraorders Gerromorpha (Andersen, 1982) and Enicocephalomorpha (Baňař and Štys, 2006; Štys and Baňař, 2007, 2013). These lineages occur among the early-diverging branches of Heteroptera, and are closely related (Knyshov et al., 2021; Song et al., 2024; Weirauch et al., 2019).

Comparable trochanteral fields have been reported in other hemipterans, including cicadellids (D’Urso and Ippolito, 1988) and aphids (Montagano and Favret, 2016), and across other insect orders (Eich et al., 2025). Examples include Phasmatodea (Hofmann and Bässler, 1982), Dictyoptera (Zill et al., 1999), Orthoptera (Mücke, 1991), Hymenoptera (Markl, 1970; Schmidt and Smith, 1987), Lepidoptera (Kent and Griffin, 1990; Kent et al., 1996), and Diptera (Dinges et al., 2021). Trochanteral fields therefore appear to be widespread among insects. CS-like structures have also been recorded on the thorax and abdomen of Enicocephalomorpha (Davranoglou et al., 2017), suggesting that the diversity and distribution of these receptors in Hemiptera remain incompletely characterised.

A comparative map of CS distribution is needed to identify candidate receptors for physiological and behavioural studies of vibroacoustic communication. Here, we survey midleg CS across a broad, but non-exhaustive, sample of Hemiptera, with particular emphasis on Heteroptera. Building on the interordinal survey of Eich et al. (2025), we address two questions: how does the number of CS vary among hemipteran lineages and leg segments, and are CS counts associated with documented vibrational communication?

We focus on the midleg because forelegs are frequently specialised for prey capture, whereas hindlegs may be modified for jumping or defence; midlegs often retain more generalised locomotor roles (Schuh and Slater, 1995). This focus provides a consistent anatomical basis for comparison, although it does not imply that midlegs are functionally equivalent across all sampled taxa. The behavioural comparison concerns documented vibrational communication; it does not directly test sensitivity to airborne sound.

## Methods

### Taxon selection and specimen sampling

Taxa were selected to represent major branches of the hemipteran phylogeny of Johnson et al. (2018), subject to specimen availability. The sample included representatives of Sternorrhyncha, Coleorrhyncha, Auchenorrhyncha and all seven heteropteran infraorders. For a summary of the examined taxa, refer to Table S1.

The non-heteropteran taxa were *Orthezia* sp. (Sternorrhyncha: Ortheziidae), *Peloridium pomponorum* (Coleorrhyncha: Peloridiidae), *Latematium* sp. (Auchenorrhyncha: Fulgoromorpha: Issidae) and *Cicada orni* (Auchenorrhyncha: Cicadomorpha: Cicadidae).

The Heteroptera comprised *Ceratocombus coleoptratus* (Dipsocoromorpha: Ceratocombidae); *Cocles* sp., provisionally designated as a new species (Enicocephalomorpha: Enicocephalidae); *Velia* sp. (Gerromorpha: Veliidae); *Anisops sardeus* (Nepomorpha: Notonectidae); an unidentified saldid from Dystos, Euboea, Greece (Leptopodomorpha: Saldidae); *Zelus renardii* (Cimicomorpha: Reduviidae: Harpactorinae); *Stenodema laevigata* (Cimicomorpha: Miridae); an unidentified mezirine, provisionally designated as a new species (Pentatomomorpha: Aradoidea: Aradidae: Mezirinae); *Codophila varia* (Pentatomomorpha: Pentatomoidea: Pentatomidae); *Lygaeus creticus* (Pentatomomorpha: Lygaeidae); and *Pyrrhocoris apterus* (Pentatomomorpha: Pyrrhocoridae).

### Preparation for scanning electron microscopy (SEM)

Legs from one side of each specimen were removed using shaped dissecting forceps. The left side was used except in *Anisops sardeus, Ceratocombus coleoptratus, Cocles* sp., *Cicada orni*, the mezirine and *Pyrrhocoris apterus*. Specimens stored in 70% ethanol were dehydrated through 80%, 90%, 95% and 100% ethanol for 1 h at each concentration, followed by a second change of 100% ethanol. They were then critical-point dried using a Tousimis Autosamdy-931 instrument for 6 cycles at 199 PSI, 20s fill, 2 min purge, 20s post-purge. Critical point was held for 2 min. Pressure was released slowly until 550 PSI, stasis time 40 min, followed by a purge of 1 min.

Following critical-point drying, specimens were immersed in 50% w/w hydrogen peroxide (H_2_O_2_) for 10, 20 or 40 min, according to specimen size, robustness and cleaning requirements. They were rinsed twice in 100% ethanol for 2 min per rinse and dried on filter paper in a desiccator. Samples were mounted on Katana holder rods, 1 cm long and 1.5 mm wide, allowing rotation through 360° using a holder based on the design of Pohl (2010), which at the same time acts as an “electron trap” thereby improving sample-to-background contrast. They were sputter-coated with palladium in a Cressington 208 HR coater for two 80-s cycles at 60mA.

### Image acquisition and sensilla mapping

High resolution overview and detailed images were acquired with a TESCAN CLARA scanning electron microscope using an Everhart–Thornley detector at an accelerating voltage of 5 kV and a reported beam current of 300 pA. Working distances ranged from approximately 9 to 35 mm, depending on specimen size and stage orientation; the longer working distances were used for overview images.

Leg surfaces were examined systematically using a field of view of approximately 100 µm (approximately 5,600× magnification), with 20–30% overlap between successive views. Selected views containing CS or candidate CS were saved.

Typically, CS are described using the nomenclature of Dinges et al. (2021). Because applying this nomenclature across taxa of different sizes can be problematic, we primarily report counts at each location and on each leg segment.

Images acquired at approximately 8,000× and 16,000× magnification were used to identify CS manually and locate them in segment-level and whole-leg overview images. Schematics were generated semi-automatically from posterior views of the midlegs using Adobe Photoshop CS5, and CS markers were placed using Adobe Illustrator CS5.

### Comparison with vibrational communication

Counts were summarised by leg segment and for the midleg as a whole. Coxae were excluded from the comparative leg totals because they were not retained in all preparations and recognisable coxal CS were found in only a subset of taxa. Confidently identified CS and candidate CS were recorded separately, with candidate counts shown in parentheses. Taxa were grouped according to whether vibrational communication had been documented or no such record was available, and the counts were displayed alongside the phylogeny of Johnson et al. (2018). The absence of a record was treated as a gap in knowledge, rather than evidence that a taxon does not communicate vibrationally.

Using the summarised CS counts per leg (without candidates), statistics were performed with Matlab2023a. It was tested whether at a significance level of α=0.05 there is a statistical difference detectable between the two groups of species with documented vibration communication (n=9) and those without documented vibration communication (n=5). After verifying that no group is normally distributed and there is no equal variance between them, a Kruskal-Wallis test was performed. The first assumption was tested with the Lilliefors test, and the second with a Levene test modified after Brown and Forsythe.

## Results

Campaniform sensilla were distributed across several midleg segments, with substantial variation among taxa (Fig. 1; Table 1). In the examined taxa other than *Orthezia* sp. (Sternorrhyncha), whose trochanter was reduced, CS occurred at two trochanteral locations. Although the segment with the highest total count varied among taxa, those trochanteral positions – anterior and posterior – contained the largest local concentrations of CS on the legs.

**Table 1:** Midleg campaniform sensillum counts. Confirmed campaniform sensilla (CS) are followed by additional candidate CS in parentheses. Both whole-leg and non-coxal totals are reported. The dash (-) stands for segment inaccessible or too damaged.

| Lineage | Taxon | Family | Coxa | Trochanter | Femur | Tibia | Tarsus | Total<br>with coxa | Total<br>without coxa |
| --- | --- | --- | --- | --- | --- | --- | --- | --- | --- |
| Pentatomomorpha | Mezirinae sp. | Aradidae | 0(1) | 8(2) | 4(9) | 2(11) | 0(1) | **14(24)** | **14(23)** |
| Pentatomomorpha | *Codophila varia* | Pentatomidae | 15(26) | 11(14) | 14(53) | 20(30) | 6(6) | **66(129)** | **51(103)** |
| Pentatomomorpha | *Pyrrhocoris apterus* | Pyrrhocoridae | 9 | 13 | 16(6) | 25(7) | 13(1) | **76(14)** | **67(14)** |
| Pentatomomorpha | *Lygaeus creticus* ‡ | Lygaeidae | 9(5) | 20(5) | 36(15) | 34(3) | 14(8) | **113(36)** | **104(31)** |
| Cimicomorpha | *Stenodema laevigata* | Miridae | 5 | 14 | 13(9) | 9(5) | 0(2) | **41(16)** | **36(16)** |
| Cimicomorpha | *Zelus renardii* | Reduviidae | 0 | 10(4) | 3(11) | 5 | 3(11) | **21(26)** | **21(26)** |
| Leptopodomorpha | Saldidae indet. | Saldidae | 3 | 15 | 4(2) | 3(2) | 1 | **26(4)** | **23(4)** |
| Nepomorpha | *Anisops sardeus* | Notonectidae | 0(2) | 7(2) | 10(10) | 12(1) | 11(2) | **40(17)** | **40(15)** |
| Gerromorpha | *Velia* sp. | Veliidae | 0 | 4(2) | 0(4) | 0 | 1 | **5(6)** | **5(6)** |
| Enicocephalomorpha | *Cocles* sp. † | Enicocephalidae | 0 | - | 4(3) | 0(2) | 1(1) | **5(6)** | **5(6)** |
| Dipsocoromorpha | *Ceratocombus coleoptratus* | Ceratocombidae | 0 | 14(1) | 2(1) | 0(5) | 1(2) | **17(9)** | **17(9)** |
| Cicadomorpha | *Cicada orni* | Cicadidae | 1(35) | 18(24) | 3(23) | 6(23) | 3(10) | **31(115)** | **30(80)** |
| Fulgoromorpha | *Latematium* sp. | Issidae | 0(1) | 20(1) | 12(1) | 13(2) | 4(1) | **49(6)** | **49(5)** |
| Coleorrhyncha | *Peloridium pomponorum* | Peloriidiidae | 0(3) | 14 | 3(1) | 0(1) | 1 | **18(5)** | **18(2)** |
| Sternorrhyncha | *Orthezia* sp. | Ortheziidae | 0(1) | 0 | 2(1) | 1(2) | 0 | **3(4)** | **3(3)** |

**Fig. 1:**
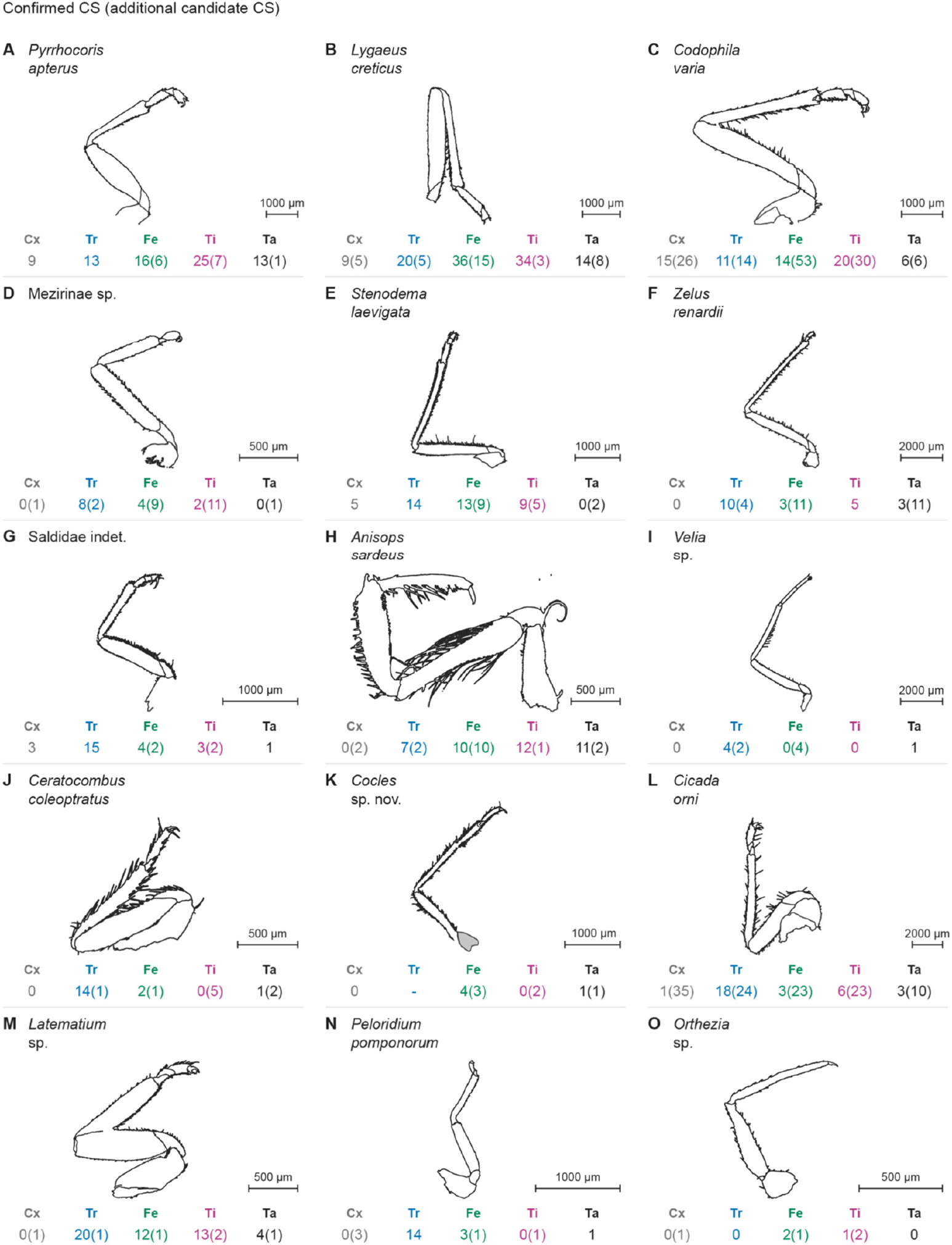
Midleg morphology and campaniform sensillum counts in the sampled Hemiptera. Schematics are shown in posterior view. Numbers outside parentheses denote confidently identified CS; those in parentheses denote additional candidate CS. Number colours identify the coxa (grey), trochanter (blue), femur (green), tibia (magenta) and tarsus (black). Scale bars explained in figure panels, corresponding to 500, 1,000 or 2,000 µm, respectively.

In *Codophila varia* and *Pyrrhocoris apterus*, the tibia bore the highest number of confidently identified CS. In *Anisops sardeus*, femoral and tibial counts were similar. The sampled representatives of Pentatomomorpha Trichophora, namely *Codophila varia, Lygaeus creticus* and *Pyrrhocoris apterus*, had particularly high overall counts relative to the sampled aradid outgroup (Table 1; Fig. 2). The trochanter also universally shows the largest number of CS in one location. And with the exception of *Anisops sardeus, Stenodema laevigata*; *Codophila varia, Lygaeus creticus, Pyrrhocoris apterus*, this is also the segment with the overall largest numbers of CS in the investigated species. Candidate CS occurred on several segments, and were numerous in some taxa, emphasising the distinction between confidently identified sensilla and structures requiring further examination.

**Figure 2:**
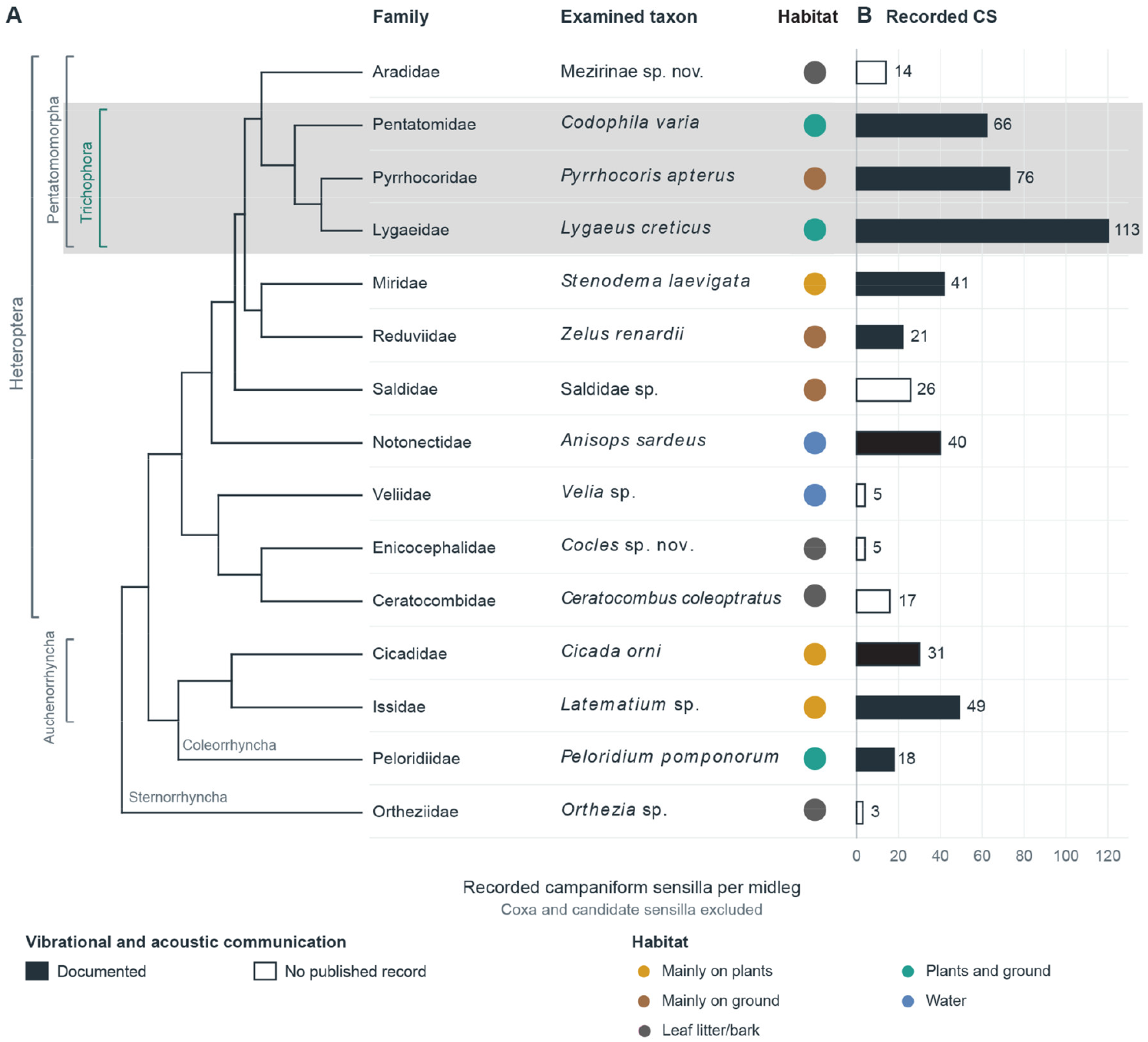
Summary of CS count, plotted in phylogenetic context. A) reproduction of the phylogenetic tree by Johnson et al., 2018 and Davranoglou and Hartung (2024). B) Species list of the present study C) number of CS found on X axis vs. species on Y axis. Legend indicates habitat type as outline, and status regarding vibrational communication as filling of the bars. Bars in black indicate experimentally demonstrated capacity for vibroacoustic communication. The data make no claim about the animals’ capacity to generate vibroacoustic signals (e.g. as part of a defensive display), as this is not always correlated with the ability to perceive such signals as shown for the acoustic signal component by Davranoglou et al. (2023).

**Fig. 3:**
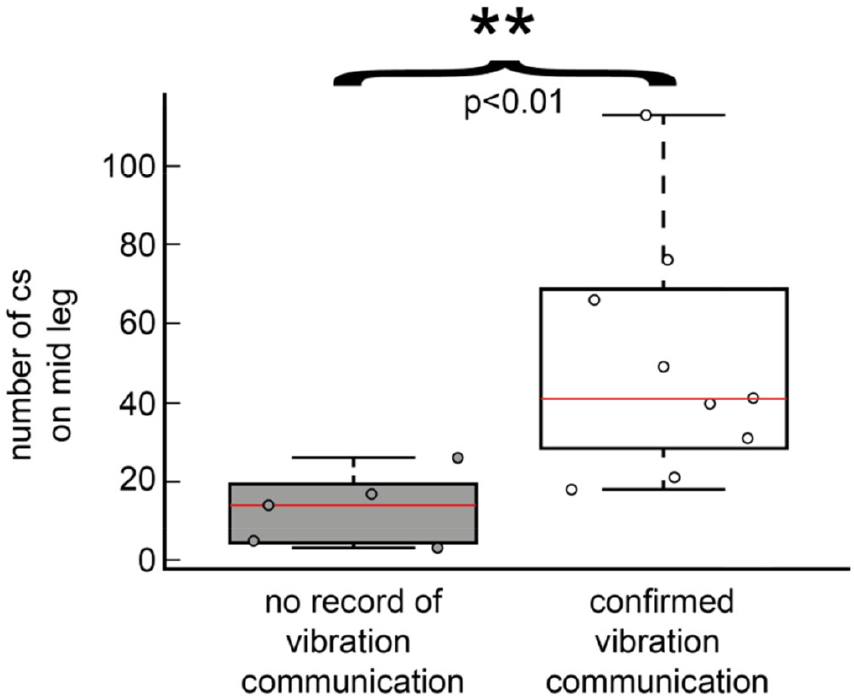
Comparison of midleg campaniform sensillum counts between taxa with documented vibrational communication and taxa for which no record was available. As the boxplot shows, there was a highly significant difference in CS number across the two groups: species with and without vibration record. Results of Kruskal-Wallis test: χ2 (df:1,12 N = 14) = 7.47, p = 0.0063. Individual dots indicate species overall count of campaniform sensilla.

Stepping back from the individual CS, we observed the number of campaniform sensilla in each leg segment and for total legs – with the exception of coxae, which did not survive sample preparation in all specimens and even then only featured CS in a small subset of species. This is shown in Fig. 2, in which we also marked known vibrational communication behaviour (as filling of the bars). Statistical results of the Kruskal-Wallis test noted in Fig. 4, show a highly significant (p<0.01, χ2 (df:1,12 N = 14) = 7.47) effect between groups with vibration communication and those without, even though sample size is limited.

## Discussion

### Comparative distribution of campaniform sensilla

Our survey documents variation in the number and distribution of midleg CS across Hemiptera. A conspicuous pattern is the high number of CS in the sampled Trichophora relative to the sampled aradid, their sister-group (Johnson et al., 2018; Song et al., 2024). Trichophora are a lineage that includes all Pentatomomorpha other than Aradidae (which lack trichobothria) (Fig. 2), and are named as such due to their possession of a large number of abdominal trichobothria, whose presence has been correlated with their capacity to generate vibroacoustic signals (Drašlar, 1973; Schaefer, 1966). However, we should note that the role of trichobothria in heteropteran vibrational communication remains unsubstantiated. The co-occurrence of these sensory features suggests a useful comparative hypothesis, but our observations do not demonstrate a quantitative correlation between CS and trichobothrial abundance.

### Interpreting the association with communication

The higher CS counts in taxa with documented vibrational communication are consistent with a possible contribution of these receptors to vibrational signal reception. However, CS are multifunctional receptors (Dickerson et al., 2021), and the observed association need not arise from the presence of vibrational communication itself. Other factors, such as shared ancestry, body or leg size, locomotor demands, habitat and other uses of mechanical information could also contribute to the observed variation in CS counts. Ergo, arranging the data alongside a phylogeny does not, by itself, account for phylogenetic non-independence. The current comparison should therefore be regarded as exploratory.

We should also note that CS number alone indicate sensitivity to vibration. The locations and orientations of CS, their mechanical coupling to the cuticle and their physiological response properties are relevant to the stimuli they encode (Dickerson et al., 2021; Lakes-Harlan and Strauß, 2014; Strauß and Stritih-Peljhan, 2022). Direct recordings and behavioural experiments are needed to determine whether the mapped sensilla respond to communication signals, locomotor forces or both. Such tests would also help distinguish a role in receiving external signals from feedback associated with an animal’s own movements.

### Morphological variation and limits of the survey

Our survey concerns externally recognisable CS; therefore, sensilla relevant to vibration that cannot be detected from the exposed surface may have been overlooked; such receptors have been documented in beetles (Ichikawa et al., 2014). Furthermore, the distinction between identified and candidate CS, together with incomplete preservation of some leg regions, also needs to be considered in our taxonomic comparison.

The variation observed here is consistent with the interordinal survey of Eich et al. (2025), while Dinges et al. (2021) documented CS variation within a species. Different sensillum configurations may provide feedback for similar mechanical tasks, and redundancy or individual variation may contribute to differences in position and number. Species-specific ecological and mechanical demands are plausible explanations, but our comparative observations do not establish their relative importance or exclude phylogenetic constraints. Demonstrating that function determines sensillum arrangement will require explicit functional tests. This includes but is not limited to electrophysiological tests.

The present survey adds an intermediate comparative scale between detailed studies of individual species and comparisons across insect orders. Further evidence of comparison of three nepomorphan species from three families was found by Nowińska et al. (2023). These comparisons concern different taxonomic levels.

## Conclusions

The distribution of midleg CS in Hemiptera combines widespread trochanteral concentrations with variation in CS number and position among leg segments. The exploratory association with documented vibrational communication provides a hypothesis for further study, rather than evidence of a causal relationship. By locating potential receptors across a broad taxonomic sample, this survey provides an anatomical basis for targeted electrophysiological and behavioural experiments. Broader sampling, explicit identification criteria and analyses that address shared ancestry and mechanical differences among taxa will be needed to evaluate the relationship between CS distribution and communication.

## Acknowledgements

Funding: Brian Saltin is supported by research grant C3NS as part of the Next Generation Networks for Neuroscience Program (DFG grant, BL1355/6-2 [436258345]). Leonidas-Romanos Davranoglou is a Leverhulme Trust Early Career Fellow, also receiving support of the John Fell OUP Fund.

Acknowledgements: Brian Saltin thanks Alexander Blanke for his support of this study.

## Notes

### Competing Interest Statement

The authors have declared no competing interest.

## References

Ai, H., Yoshida, A., Yokohari, F., 2010. Vibration receptive sensilla on the wing margins of the silkworm moth Bombyx mori. Journal of Insect Physiology 56, 236–246. 10.1016/j.jinsphys.2009.10.007

Akay, T., Bässler, U., Gerharz, P., Büschges, A., 2001. The role of sensory signals from the insect coxa-trochanteral joint in controlling motor activity of the femur-tibia joint. Journal of Neurophysiology 85, 594–604.

Andersen, N.M., 1982. The semiaquatic bugs (Hemiptera, Gerromorpha): phylogeny, adaptations, biogeography and classification. Entomonograph 3, 1–455.

Baňař, P., Štys, P., 2006. A new species of Pseudohenschiella (Heteroptera: Enicocephalidae) from Madagascar. Acta Entomologica Musei Nationalis Pragae 46, 7–14.

Brożek, J., Chłond, D., 2010. Morphology, arrangement and classification of sensilla on the apical segment of labium in Peiratinae (Hemiptera: Heteroptera: Reduviidae). Zootaxa 2476, 39–52. 10.11646/zootaxa.2476.1.5

Büschges, A., Akay, T., Gabriel, J.P., Schmidt, J., 2008. Organizing network action for locomotion: Insights from studying insect walking. Brain Research Reviews 57, 162–171. 10.1016/j.brainresrev.2007.06.028

Cokl, A., Amon, T., 1980. Vibratory Interneurons in the Central Nervous System of Nezara viridula L. (Pentatomidae, Heteroptera). Journal of Comparative Physiology 139, 87–95.

Davranoglou, L.-R., Baňař, P., Schlepütz, C.M., Mortimer, B., Taylor, G.K., 2017. The pregenital abdomen of Enicocephalomorpha and morphological evidence for different modes of communication at the dawn of heteropteran evolution. Arthropod Structure & Development 46, 843–868. 10.1016/j.asd.2017.08.006

Davranoglou, L. R., Hartung, V., 2024. Moss bugs shed light on the evolution of complex bioacoustic systems. Plos one, 19, 2, e0298174. 10.1371/journal.pone.0298174

Davranoglou, L. R., Taylor, G. K., & Mortimer, B., 2023. Sexual selection and predation drive the repeated evolution of stridulation in Heteroptera and other arthropods. Biological Reviews, 98(3), 942–981. DOI: 10.1111/brv.12938

Dickerson, B.H., Fox, J.L., Sponberg, S., 2021. Functional diversity from generic encoding in insect campaniform sensilla. Current Opinion in Physiology 19, 194–203.

Dickinson, M.H., 1999. Haltere-mediated equilibrium reflexes of the fruit fly, Drosophila melanogaster. Philosophical Transactions of the Royal Society of London. Series B, Biological Sciences 354, 903–916. 10.1098/rstb.1999.0442

Dinges, G.F., Chockley, A.S., Bockemühl, T., Ito, K., Blanke, A., Büschges, A., 2021. Location and arrangement of campaniform sensilla in Drosophila melanogaster. Journal of Comparative Neurology 529, 905–925. 10.1002/cne.24987

Drašlar, K., 1973. Functional properties of trichobothria in the bug Pyrrhocoris apterus (L.). Journal of comparative physiology, 84, 2, 175-184.

D’Urso, V., Ippolito, S., 1988. Proprioceptor organs in Cicadella viridis (L.) (Homoptera : Cicadellidae). International Journal of Insect Morphology and Embryology 17, 381–391. 10.1016/0020-7322(88)90018-9

Eich, R.G.F., Lemmen, J.I., Saltin, B., Blanke, A., 2025. An overview of campaniform sensilla arrangement and location on hexapod legs. Arthropod Structure & Development 88, 101477. 10.1016/j.asd.2025.101477

Harris, C.M., Szczecinski, N.S., Büschges, A., Zill, S.N., 2022. Sensory signals of unloading in insects are tuned to distinguish leg slipping from load variations in gait: experimental and modeling studies. Journal of Neurophysiology 128, 790–807. 10.1152/jn.00285.2022

Hofmann, T., Bässler, U., 1982. Anatomy and physiology of trochanteral campaniform sensilla in the stick insect, Cuniculina impigra. Physiological Entomology 7, 413–426.

Ichikawa, T., Toh, Y., Ohkubo, K., Nishino, H., 2014. Microscopic analysis of mechanosensory system monitoring the dynamic claw actions in the tenebrionid beetle Zophobas atratus. Zoomorphology 133, 273–284. 10.1007/s00435-014-0225-8

Johnson, K.P., Dietrich, C.H., Friedrich, F., Beutel, R.G., Wipfler, B., Peters, R.S., Allen, J.M., Petersen, M., Donath, A., Walden, K.K.O., Kozlov, A.M., Podsiadlowski, L., Mayer, C., Meusemann, K., Vasilikopoulos, A., Waterhouse, R.M., Cameron, S.L., Weirauch, C., Swanson, D.R., Percy, D.M., Hardy, N.B., Terry, I., Liu, S., Zhou, X., Misof, B., Robertson, H.M., Yoshizawa, K., 2018. Phylogenomics and the evolution of hemipteroid insects. Proceedings of the National Academy of Sciences 115, 12775–12780. 10.1073/pnas.1815820115

Keil, T.A., 1997. Functional morphology of insect mechanoreceptors. Microscopy Research and Technique 39, 506–531. 10.1002/(SICI)1097-0029(19971215)39:6%3C506::AID-JEMT5%3E3.0.CO;2-B

Kent, K.S., Fjeld, C.C., Anderson, R., 1996. Leg proprioceptors of the tobacco hornworm, Manduca sexta: organization of central projections at larval and adult stages. Microscopy Research and Technique 35, 265–284. 10.1002/(SICI)1097-0029(19961015)35:3%3C265::AID-JEMT6%3E3.0.CO;2-M

Kent, K.S., Griffin, L.M., 1990. Sensory organs of the thoracic legs of the moth Manduca sexta. Cell and Tissue Research 259, 209–223. 10.1007/BF00318442

Knyshov, A., Weirauch, C., Hoey-Chamberlain, R., 2021. Phylogenetic relationships and revised classification of the true bug infraorder Dipsocoromorpha (Insecta: Hemiptera: Heteroptera). Cladistics 37, 248–275. 10.1111/cla.12435

Kühne, R., 1982. Neurophysiology of the vibration sense in locusts and bushcrickets: Response characteristics of single receptor units. Journal of Insect Physiology 28, 155–163. 10.1016/0022-1910(82)90123-8

Lakes-Harlan, R., Strauß, J., 2014. Functional Morphology and Evolutionary Diversity of Vibration Receptors in Insects, in: Cocroft, R.B., Gogala, M., Hill, P.S.M., Wessel, A. (Eds.), Studying Vibrational Communication. Springer, Berlin, Heidelberg, pp. 277–302. 10.1007/978-3-662-43607-3_14

Markl, H., 1970. Die Verständigung durch Stridulationssignale bei Blattschneiderameisen. Zeitschrift für vergleichende Physiologie 69, 6–37.

McIver, S.B., 1975. Structure of cuticular mechanoreceptors of arthropods. Annual Review of Entomology 20, 381–397.

Mendes, C. S., Rajendren, S. V., Bartos, I., Márka, S., Mann, R. S., 2014. Kinematic responses to changes in walking orientation and gravitational load in Drosophila melanogaster. PloS one, 9, 10, e109204.

Montagano, L., Favret, C., 2016. The Distribution of Campaniform Sensilla on the Appendages of Mindarus Species (Hemiptera: Aphididae). Entomological News 126, 196–203. 10.3157/021.126.0305

Mücke, A., 1991. Innervation pattern and sensory supply of the midleg of Schistocerca gregaria (Insecta, Orthopteroidea). Zoomorphology 110, 175–187.

Nowińska, A., Franielczyk-Pietyra, B., Polhemus, D.A., 2023. The Leg Sensilla of Insects from Different Habitats—Comparison of Strictly Aquatic and Riparian Bugs (Corixidae, Ochteridae, Gelastocoridae: Nepomorpha: Insecta: Heteroptera). Insects 14, 441. 10.3390/insects14050441

Parveen, S., Ahmad, A., Brożek, J., Ramamurthy, V.V., 2015. Morphological diversity of the labial sensilla of phytophagous and predatory Pentatomidae (Hemiptera: Heteroptera), with reference to their possible functions. Zootaxa 4039, 359–372. 10.11646/zootaxa.4039.2.9

Pohl, H., 2010. A scanning electron microscopy specimen holder for viewing different angles of a single specimen. Microscopy Research and Technique 73, 1073–1076.

Pringle, J.W.S., 1948. The gyroscopic mechanism of the halteres of Diptera. Philosophical Transactions of the Royal Society of London. Series B, Biological Sciences 233, 347–384. 10.1098/rstb.1948.0007

Schaefer, C. W., 1966. Some notes on heteropteran trichobothria. The Great Lakes Entomologist, 1, 85–90.

Schmidt, J.M., Smith, J.J.B., 1987. The external sensory morphology of the legs and hairplate system of female Trichogramma minutum Riley (Hymenoptera: Trichogrammatidae). Proceedings of the Royal Society of London. Series B. Biological Sciences 232, 323–366.

Schuh, R.T., Slater, J.A., 1995. True bugs of the world: Hemiptera, Heteroptera classification and natural history. Cornell University Press, Ithaca.

Simmons, P.J., 2024. Scaling of sense organs that control flight: Size and sensory cell number of dipteran fly halteres. Journal of Zoology 322, 12–23. 10.1111/jzo.13117

Smith, D.S., 1969. The fine structure of haltere sensilla in the blowfly Calliphora erythrocephala (Meig.), with scanning electron microscopic observations on the haltere surface. Tissue and Cell 1, 443–484. 10.1016/S0040-8166(69)80016-9

Song, N., Wang, M.-M., Huang, W.-C., Wu, Z.-Y., Shao, R., Yin, X.-M., 2024. Phylogeny and evolution of hemipteran insects based on expanded genomic and transcriptomic data. BMC Biology 22, 190. 10.1186/s12915-024-01991-1

Strauß, J., 2020. Neuronal Innervation of the Subgenual Organ Complex and the Tibial Campaniform Sensilla in the Stick Insect Midleg. Insects 11, 40. 10.3390/insects11010040

Strauß, J., Stritih-Peljhan, N., 2022. Vibration detection in arthropods: Signal transfer, biomechanics and sensory adaptations. Arthropod Structure & Development 68, 101167.

Štys, P., Baňař, P., 2013. Eastern Arc Mountains in Tanzania: Hic sunt Aenictopecheidae. The first genus and species of Afrotropical Aenictopecheidae (Hemiptera: Heteroptera: Enicocephalomorpha). European Journal of Entomology 110, 677–688.

Štys, P., Baňař, P., 2007. The first species of Systelloderes (Hemiptera: Heteroptera: Enicocephalidae) from New Caledonia. Acta Entomologica Musei Nationalis Pragae 47, 3–15.

Taszakowski, A., Nowińska, A., Brożek, J., 2019. Morphological study of the labial sensilla in Nabidae (Hemiptera: Heteroptera: Cimicomorpha). Zoomorphology 138, 483–492. 10.1007/s00435-019-00455-3

Wang, Y., Dai, W., 2017. Fine structure of mouthparts and feeding performance of Pyrrhocoris sibiricus Kuschakevich with remarks on the specialization of sensilla and stylets for seed feeding. PLoS ONE 12, e0177209. 10.1371/journal.pone.0177209

Weirauch, C., Schuh, R.T., Cassis, G., Wheeler, W.C., 2019. Revisiting habitat and lifestyle transitions in Heteroptera (Insecta: Hemiptera): insights from a combined morphological and molecular phylogeny. Cladistics 35, 67–105. 10.1111/cla.12233

Yarger, A.M., Fox, J.L., 2018. Single mechanosensory neurons encode lateral displacements using precise spike timing and thresholds. Proceedings of the Royal Society B: Biological Sciences 285, 20181759. 10.1098/rspb.2018.1759

Zill, S., Schmitz, J., Büschges, A., 2004. Load sensing and control of posture and locomotion. Arthropod Structure & Development 33, 273–286. 10.1016/j.asd.2004.05.005

Zill, S.N., Büschges, A., Schmitz, J., 2011. Encoding of force increases and decreases by tibial campaniform sensilla in the stick insect, Carausius morosus. Journal of Comparative Physiology A 197, 851–867. 10.1007/s00359-011-0647-4

Zill, S.N., Chaudhry, S., Exter, A., Büschges, A., Schmitz, J., 2014. Positive force feedback in development of substrate grip in the stick insect tarsus. Arthropod Structure & Development 43, 441–455.

Zill, S.N., Keller, B.R., Chaudhry, S., Duke, E.R., Neff, D., Quinn, R., Flannigan, C., 2010. Detecting substrate engagement: responses of tarsal campaniform sensilla in cockroaches. Journal of Comparative Physiology A 196, 407–420.

Zill, S.N., Keller, B.R., Duke, E.R., 2009. Sensory signals of unloading in one leg follow stance onset in another leg: transfer of load and emergent coordination in cockroach walking. Journal of Neurophysiology 101, 2297–2304. 10.1152/jn.00056.2009

Zill, S.N., Moran, D.T., 1981. The exoskeleton and insect proprioception. I. Responses of tibial campaniform sensilla to external and muscle-generated forces in the American cockroach, Periplaneta americana. Journal of Experimental Biology 91, 1–24.

Zill, S.N., Ridgel, A.L., DiCaprio, R.A., Frazier, S.F., 1999. Load signalling by cockroach trochanteral campaniform sensilla. Brain Research 822, 271–275.

